# Pangenome- and genome-based taxonomic classification inference for the marine bacterial strain KMM 296 producing a highly active PhoA alkaline phosphatase and closely related *Cobetia* species

**DOI:** 10.1101/2023.11.15.567287

**Authors:** O.I. Nedashkovskaya, L.A. Balabanova, N. Y. Otstavnykh, N.V. Zhukova, A.V. Seitkalieva, Yu.A. Noskova, L.A Tekutyeva

## Abstract

A strictly aerobic, Gram-stain-negative, rod-shaped and motile bacterium, designated strain KMM 296, isolated from the coelomic fluid of mussel *Crenomytilus grayanus*, was investigated in details due to its ability to produce a highly active alkaline phosphatase of the structural family PhoA. A previous taxonomic study placed the strain to the species *Cobetia marina*, a member of the family *Halomonadaceae* of the class *Gammaproteobacteria*. However, the comprehensive phylogenetic analysis based on 16S rRNA gene sequencing revealed that the strain KMM 296 is most closely related to *Cobetia amphilecti* NRIC 815^T^ with the 16S rRNA gene sequence similarity of 100%. The mussel isolate grew with 0.5-19% NaCl and at 4 - 42°C and hydrolysed Tweens 20 and 40, and L-tyrosine. The DNA G+C content was 62.5 mol%. The prevalent fatty acids were C_18:1_ ω7*c*, C_12:0_ 3-OH, C_18:1_ ω7*c*, C_12:0_ and C_17:0_ cyclo. The polar lipid profile was characterized by the presence of phosphatidylethanolamine, phosphatidylglycerol, phosphatidic acid, and unidentified aminolipid, phospholipid, and lipids. The major respiratory quinone was Q-8. According to phylogenetic evidence and similarity in the chemotaxonomic and genotypic properties of the mussel isolate and its nearest neighbors, the strain KMM 296 represents a member of the species *C. amphilecti*. A comparative analysis of the type strains genomes of the species *C. amphilecti* and *C. litoralis* showed that they belong to a single species. In addition, a high similarity of the genome sequences of *C. pacifica* NRIC 813^T^ and *C. marina* LMG 2217^T^ also allows suggesting the affiliation of these two species to one species. Based on the rules of priority, *C. litoralis* should be reclassified as a later heterotypic synonym of *C. amphilecti*, and *C. pacifica* is a later heterotypic synonym of *C. marina*. The emended descriptions of the species *C. amphilecti* and *C. marina* are also proposed.

## Introduction

The genus *Cobetia* was created by Arahal et al. [1] to reclassify the species *Halomonas marina*. At the time of writing, the genus *Cobetia* comprises 5 validly published species, including *C. marina* as the type species, *C. amphilecti, C. crustatorum, C. litoralis*, and *C. pacifica* isolated from different marine environments [1-3]. The genus accommodates Gram-stain-negative, aerobic, heterotrophic, halophilic and rod-shaped bacteria, which can motile by means of a single polar flagellum and/or two to seven lateral flagella [1-3]. Earlier, in a course of survey of the *Halomonas*-like bacteria inhabitant different areas of NW Pacifica, the strain KMM 296 was isolated from the coelomic fluid of mussel *C. grayanus*, collected from the Sea of Japan, and identified as a representative of the species *Cobetia marina* (formerly *Deleya marina*) [4, 5]. Later, the results of phylogenetic analysis based on the 16S rRNA gene sequences revealed the closest relationship of the strain KMM 296 to the strain *C. amphilecti* NRIC 815^T^ with 100% of their sequence similarity. It should be noted, that the strain KMM 296 genome was sequenced due to its ability to produce a high-active alkaline phosphatase of the PhoA family (CmAP) and deposited to the GenBank database under the accession number NZ_JQJA00000000.1 [6-9]. In the present study, we have clarified the taxonomic position of strain KMM 296 as a member of the species *C. amphilecti* and specified the description of species *C. amphilecti* based on the results of phylogenetic analysis, phenotypic characterization and experiments on the DNA-DNA hybridization between the strains KMM 296 and *C. amphilecti* NRIC 815^T^. In addition, a comparison of the genomic sequences and phenotypic characteristics suggested that the species *C. amphilecti* and *C. litoralis* belong to a single species. Moreover, based on the data obtained in this study, we proposed the species *C. marina* and *C. pacifica* to be considered the representatives of a single species. In accordance with the rules of priority, *C. litoralis* should be reclassified as a later heterotypic synonym of *C. amphilecti*, and *C. pacifica* is a later heterotypic synonym of *C. marina*. The emended descriptions of the species *C. amphilecti* and *C. marina* are also proposed.

## Methods

### Strain cultivation

Strain KMM 296 was obtained from Collection of marine microorganisms (KMM) at the G.B. Elyakov Pacific Institute of Bioorganic Chemistry FEB RAS (Vladivostok, Russia) and cultivated at 28°C on Marine agar (Difco) and stored at –80°C in Marine broth (Difco) supplemented with 20% (v/v) glycerol. The type strains *C. amphilecti* NRIC 815^T^ (= KMM 1561^T^), *C. litoralis* NRIC 814^T^ (=KMM 3880^T^), *C. pacifica* NRIC 813^T^ (= KMM 3879^T^), and *C. marina* LMG 2217^T^ were kindly provided us by the NODAI Culture Collection Center (NRIC, Tokyo University of Agriculture, Tokyo, Japan) and the Belgian Coordinated Collection of Microorganisms (BCCM, Ghent University, Ghent, Belgium), respectively, and used as the reference strains for comparative taxonomic analysis.

### Morphological, biochemical, and physiological characterization

The physiological, morphological and biochemical properties of strain KMM 296 were studied using the standard methods. The novel isolate was also examined in the API 20E, API 20NE, API 50 CH, API 32 ID GN, and API ZYM galleries (bioMérieux, France) according to the manufacturer’s instructions, except that the galleries were incubated at 28 ºC. Gram staining was performed as recommended by [10]. Oxidative or fermentative utilization of glucose was determined on Hugh and Leifson’s medium modified for marine bacteria [11]. Catalase activity was tested by addition of 3 % (v/v) H_2_O_2_ solution to a bacterial colony and observation for the appearance of gas. Oxidase activity was determined by using tetramethyl-*p*-phenylenediamine. Degradation of agar, starch, casein, gelatin, chitin, DNA, and urea and production of acid from carbohydrates, hydrolysis of Tween 80, nitrate reduction, production of hydrogen sulphide, acetoin (Voges-Proskauer reaction), and indole were tested according to standard methods [10]. The temperature range for growth was assessed on marine agar (MA). Tolerance to NaCl was assessed in medium containing 5 g Bacto Peptone (Difco), 2 g Bacto Yeast Extract (Difco), 1 g glucose, 0.02 g KH_2_PO_4_ and 0.05 g MgSO_4_·7H_2_O per liter of distilled water with 0, 0.5, 1.0, 1.5, 2.0, 2.5, 3, 4, 5, 6, 8, 10, 12. 15, 17, 19, and 20% (w/v) of NaCl. Susceptibility to antibiotics was examined by the routine disc diffusion plate method. Discs were impregnated with the following antibiotics: ampicillin (10 μg), benzylpenicillin (10U), carbenicillin (100 μg), cefalexin (30 μg), cefazolin (30 μg), chloramphenicol (30 μg), erythromycin (15 μg), gentamicin (10 μg), kanamycin (30 μg), lincomycin (15 μg), nalidixic acid (30 μg), neomycin (30 μg), ofloxacin (5 μg), oleandomycin (15 μg), oxacillin (10 μg), polymyxin B (300 U), rifampicin (5 μg), streptomycin (30 μg), tetracycline (5 μg) and vancomycin (30 μg).

### Whole cell fatty acid, polar lipid and respiratory quinone composition

For the comparative whole-cell fatty acid and polar lipid analysis, the strains KMM 296 and *C. amphilecti* NRIC 815T were grown under optimal physiological conditions for both strains (at 30 °C for 24 h on MA). Cellular fatty acid methyl esters (FAMEs) were prepared according to the methods described by Sasser [12] using the standard protocol of Sherlock Microbial Identification System (version 6.0, MIDI) and analysed using a GC-21A chromatograph (Shimadzu) equipped with a fused-silica capillary column (30 m × 0.25 mm) coated with Supercowax-10 and SPB-5 phases (Supelco) at 210°C. FAMEs were identified by using equivalent chain-length measurements and comparing the retention times to those of authentic standards. The polar lipids of the strains studied were extracted using the chloroform/methanol extraction method of Bligh and Dyer [13]. Two-dimensional TLC of polar lipids was carried out on silica gel 60 F254 (10 × 10 cm; Merck) using chloroform/methanol/water (65: 25: 4, by vol.) in the first dimension and chloroform/methanol/acetic acid/water (80: 12: 15: 4, by vol.) in the second dimension [14]. The spray reagents used to reveal the spots was molybdophosphoric acid. Isoprenoid quinones were extracted with chloroform/methanol (2:1, v/v) and purified by TLC using a mixture of *n*-hexane and diethyl ether (85:15, v/v) as the solvent. Isoprenoid quinone composition of the strain KMM 296 was characterized by HPLC (Shimadzu LC-10A) using a reversed-phase type Supelcosil LC-18 column (15 cm × 4.6 mm) and acetonitrile/2-propanol (65:35, v/v) as a mobile phase at a flow rate of 0.5 ml min-1 as described previously [15].

### 16S rRNA gene sequencing and analysis

DNA was extracted from 0.1-0.2 g of the bacterial cells (wet weight), using an extraction protocol of Sambrook and Russell [16]. PCR was carried out using the universal oligonucleotide primers 11F (5′-GTTTGATCMTGGCTCAG-3’) and 1492R (5′-TACGGYTACCTTGTTACGACTT-3’) as described by Weisburg [17]; and the GeneAmp PCR System 9700 (Applied Biosystems Inc.). PCR amplicons were used as templates for sequencing amplification using a BigDye Terminator version 3.1 Cycle sequencing kit (Applied Biosystems). The purified sequencing products were analyzed by electrophoresis on a 50 cm capillary array of an ABI Prism 3130 DNA sequencer and the sequence was assembled with SeqScape version 2.6 (Applied Biosystems). The sequences obtained were deposited in NCBI GenBank under the succession numbers presented by Noskova et al. [18] and analyzed against the referent phylotypes, based on the type strains 16S rRNA gene sequences and whole-genome assemblies in the EzBioCloud database [19].

### Whole-genome shotgun sequencing and phylogenetic analysis

The genomic DNA was obtained from bacterial cultures of eight *Cobetia* strains *C. amphilecti* NRIC 0815^T^, *C. litoralis* NRIC 0814^T^, *C. pacifica* NRIC 0813^T^, *Cobetia* sp. 1AS1, *Cobetia* sp. 2AS, *Cobetia* sp. 3AK, *Cobetia* sp. 10Alg 146, and *Cobetia* sp. 29-18-1 using the NucleoSpin Microbial DNA Mini kit (Macherey-Nagel, Düren, Germany), following the manufacturer’s instructions. Whole-genome shotgun sequencing was carried out on an Illumina MiSeq platform using Nextera DNA Flex kits (Illumina, San Diego, CA, USA) with a 150-bp paired-end sequencing kit (Illumina, San Diego, CA, USA). The sequence quality was assessed via FastQC version 0.11.8 [FastQC. Available online: http://www.bioinformatics.babraham.ac.uk/projects/fastqc/ (accessed on 28 December 2018).], and reads were trimmed using Trimmomatic version 0.38 [20]. Filtered reads were assembled *de novo* with SPAdes version 3.15.3 [21]. The draft genomes of the strains *C. amphilecti* NRIC 0815^T^, *C. litoralis* NRIC 0814^T^, *C. pacifica* NRIC 0813^T^, *Cobetia* sp. 1AS1, *Cobetia* sp. 2AS, *Cobetia* sp. 3AK, *Cobetia* sp. 10Alg 146, and *Cobetia* sp. 29-18-1 were annotated using NCBI Prokaryotic Genome Annotation Pipeline (PGAP) [22] and deposited in GenBank/EMBL/DDBJ under the accession numbers JASCSA000000000, JARWKV000000000, JASCSB000000000, JARWKU000000000, JARWKQ000000000, JARWKR000000000, JARWKT000000000, and JARWKS000000000, respectively.

All publicly available *Cobetia* genomes were retrieved from Reference sequence (RefSeq) database at NCBI using ncbi-genome-download version 0.3.0 (https://github.com/kblin/ncbi-genome-download, accessed on 16 March 2023, n=28). The accession numbers for the genomes used in this study are listed in Table 1.

**Table 1.**
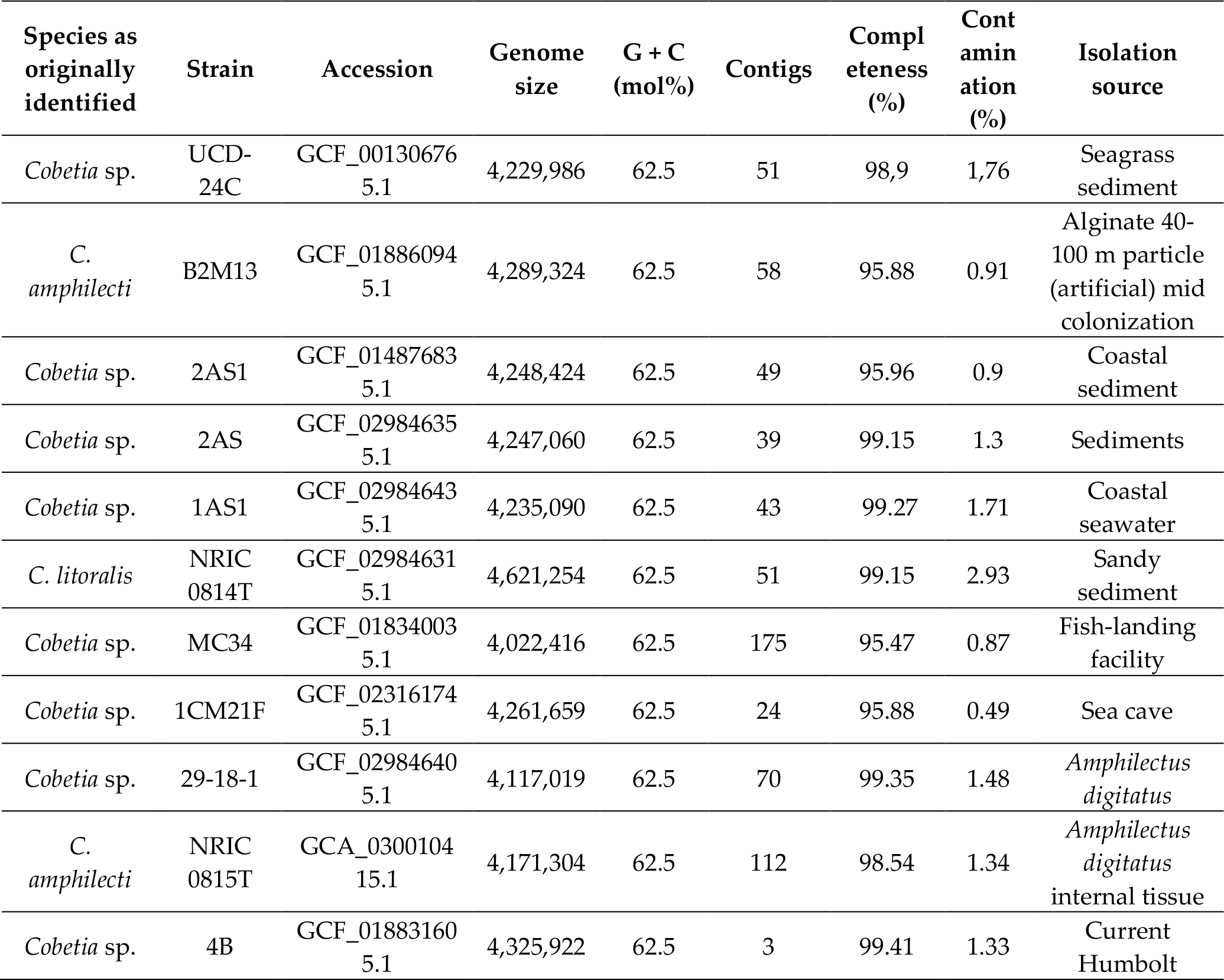
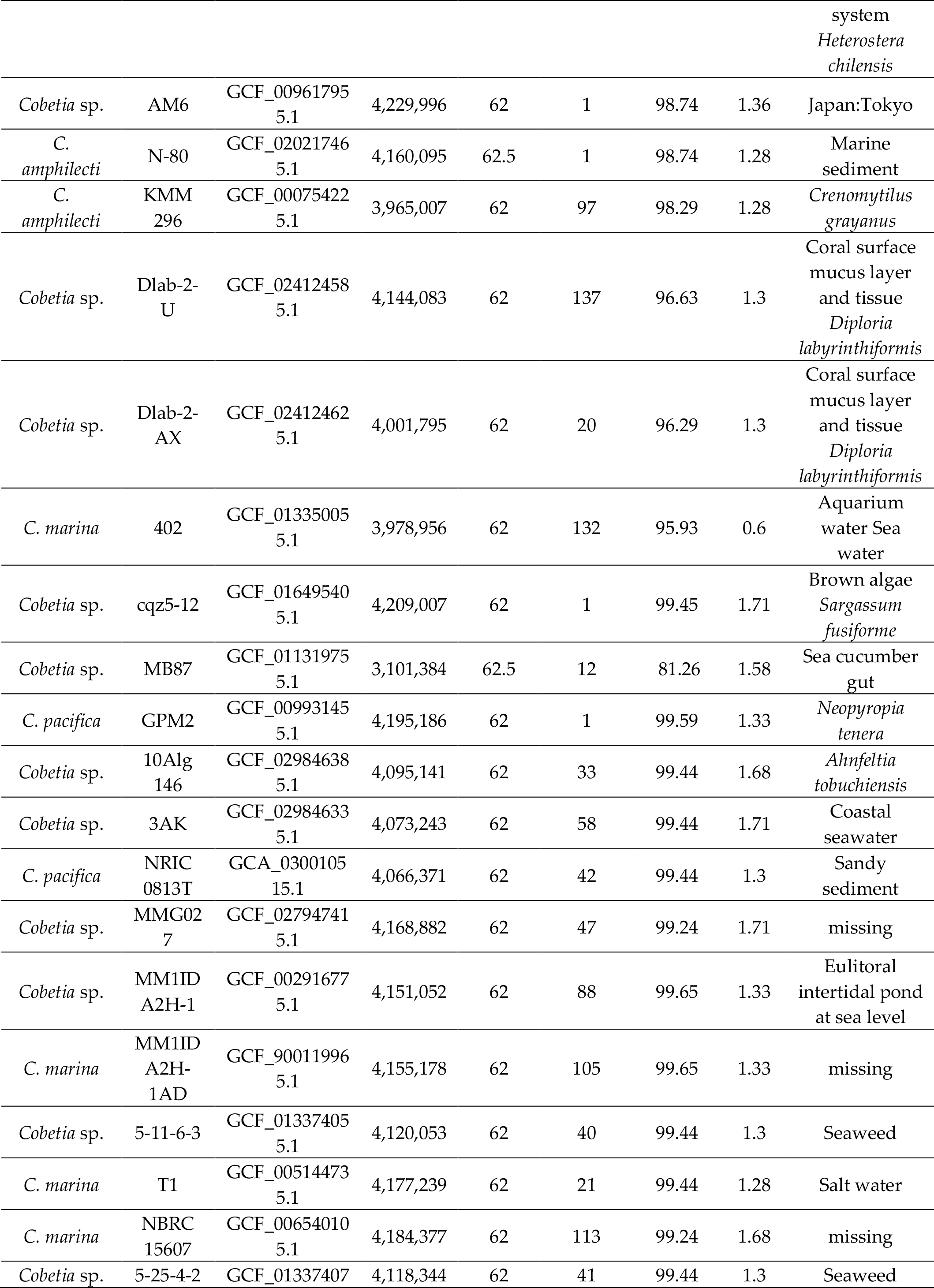
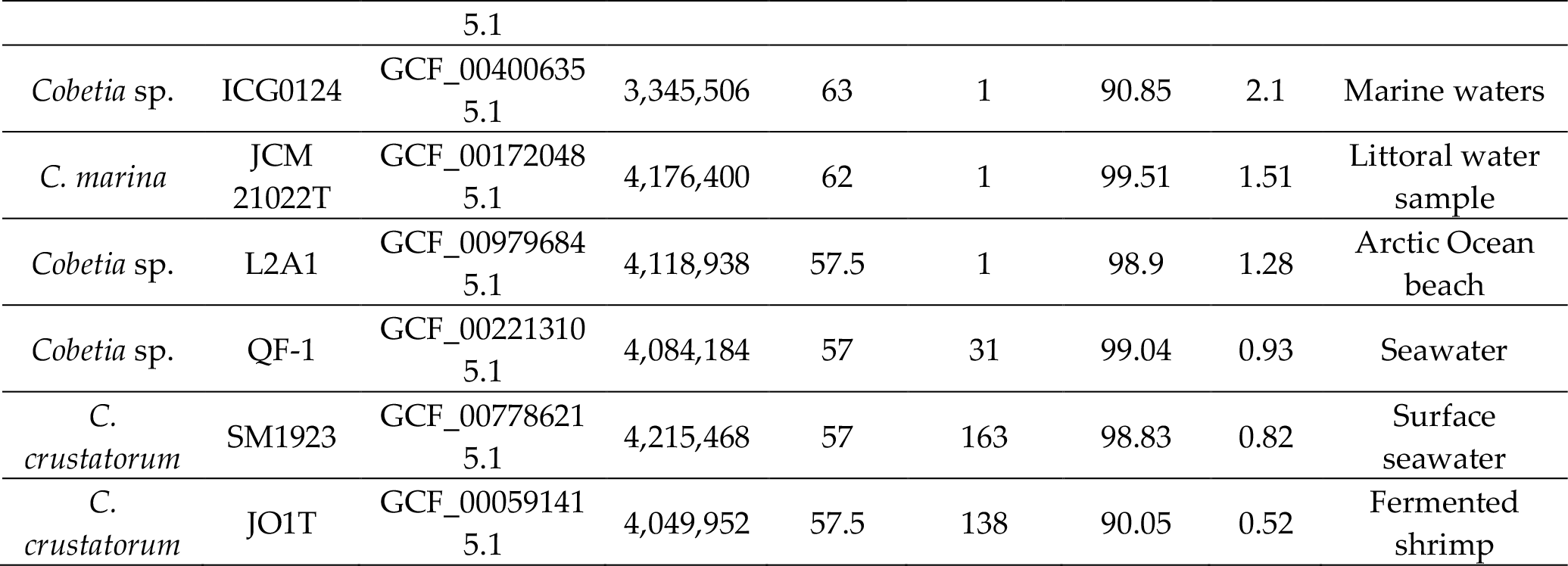
The accession numbers and general attributes of the *Cobetia* spp. genomes used in this study.

The pan-genome for these 36 *Cobetia* strains was reconstructed using the microbial pangenomics workflow in Anvi’o version 7.1 (minbit=0.5; mcl-inflation=2; min-occurrence=1) [23]. The genomes were organized based on the distribution of gene clusters using MCL algorithm (Distance: Euclidean; linkage: Ward). For the Average Nucleotide Identity (ANIm) calculations, we used the program ‘anvi-compute-genome-similarity’ with ‘--program pyANI’ flag. Amino Acid Identity (AAI) and in silico DNA–DNA hybridization (dDDH) values between the strains were calculated with the online server ANI/AAI-Matrix [24] and at the TYGS platform (formula d4), respectively [25]. To infer a phylogenetic tree of the genus *Cobetia*, 1432 single-copy core gene sequences for each strain were extracted from the pan-genome and concatenated (composite length of 465,701 bp) using program ‘anvi-get-sequences-for-gene-clusters’ with ‘--concatenate-gene-clusters’ flag. Resulted FASTA files were cleaned up by removing nucleotide positions that were gap characters in more than 50% of the sequences using trimAl version 1.4.1 [26]. A core genome phylogeny was reconstructed with IQ-TREE version 2.2.0.3 under the WAG model with non-parametric bootstrapping using 100 replicates [27]. The pangenome and core genome modelling were estimated with PanGP v.1.0.1 using a power-law regression model based on Heap’s law and exponential regression, respectively, as described by Tettelin et al [28].

Fonts and sizes in all figures were edited manually in Adobe Photoshop CC 2018 for better visualization.

## Results and Discussion

### Morphological, biochemical, and physiological characterization

The strain KMM 296 was shown to be a strictly aerobic, heterotrophic, Gram-stain-negative and motile bacterium, which formed slightly yellow-colored colony on marine agar and required NaCl or seawater for growth. The mussel isolate was positive for cytochrome oxidase and catalase and did not hydrolyse agar, casein, gelatin, starch, Tween 80, DNA, urea, and chitin (Table 2).

**Table 2.**
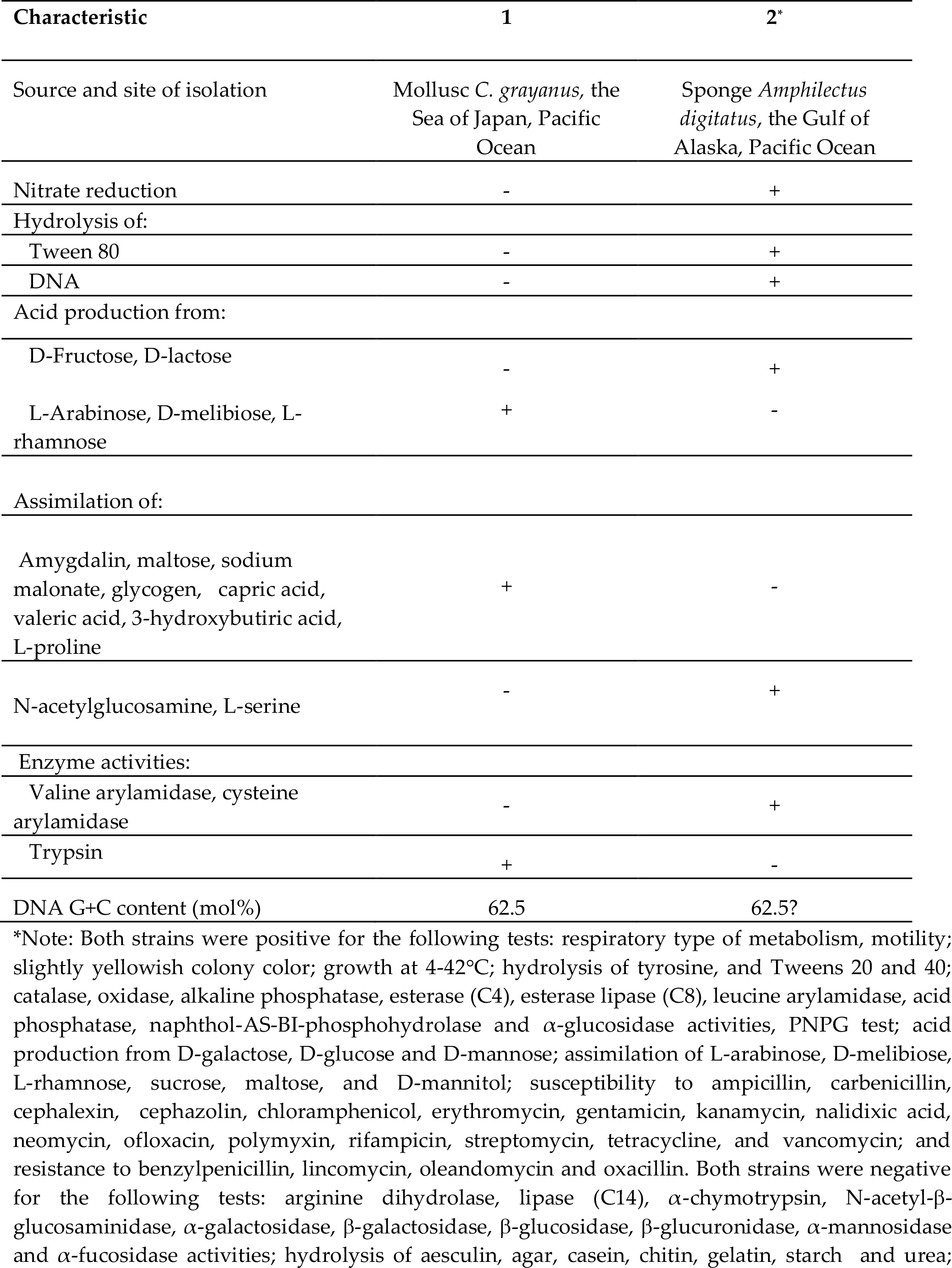
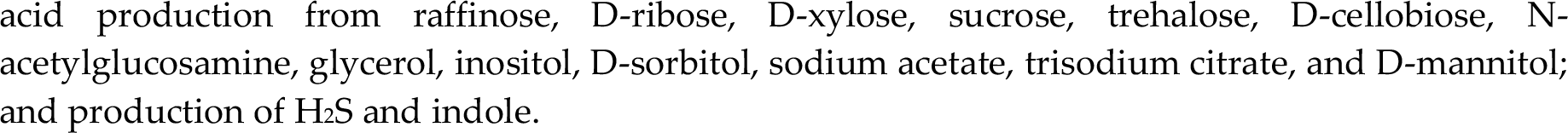
Different characteristics of the strain KMM 296 (1) and its closest relative, *Cobetia amphilecti* NRIC 815^T^ (2)*

| Characteristic | 1 | 2* |
| --- | --- | --- |
| Source and site of isolation | Mollusc *C. grayanus*, the Sea of Japan, Pacific Ocean | Sponge *Amphilectus digitatus*, the Gulf of Alaska, Pacific Ocean |
| Nitrate reduction | - | + |
| Hydrolysis of: |  |  |
| Tween 80 | - | + |
| DNA | - | + |
| Acid production from: |  |  |
| D-Fructose, D-lactose | - | + |
| L-Arabinose, D-melibiose, L-rhamnose | + | - |
| Assimilation of: |  |  |
| Amygdalin, maltose, sodium malonate, glycogen, capric acid, valeric acid, 3-hydroxybutiric acid, L-proline | + | - |
| N-acetylglucosamine, L-serine | - | + |
| Enzyme activities: |  |  |
| Valine arylamidase, cysteine arylamidase | - | + |
| Trypsin | + | - |
| DNA G+C content (mol%) | 62.5 | 62.5? |

\*Note: Both strains were positive for the following tests: respiratory type of metabolism, motility; slightly yellowish colony color; growth at 4-42°C; hydrolysis of tyrosine, and Tweens 20 and 40; catalase, oxidase, alkaline phosphatase, esterase (C4), esterase lipase (C8), leucine arylamidase, acid phosphatase, naphthol-AS-BI-phosphohydrolase and α-glucosidase activities, PNPG test; acid production from D-galactose, D-glucose and D-mannose; assimilation of L-arabinose, D-melibiose, L-rhamnose, sucrose, maltose, and D-mannitol; susceptibility to ampicillin, carbenicillin, cephalixin, cephalozin, chloramphenicol, erythromycin, gentamicin, kanamycin, nalidixic acid, neomycin, ofloxacin, polymyxin, rifampicin, streptomycin, tetracycline, and vancomycin; and resistance to benzylpenicillin, lincomycin, oleandomycin and oxacillin. Both strains were negative for the following tests: arginine dihydrolase, lipase (C14), α-chymotrypsin, N-acetyl-β-glucosaminidase, α-galactosidase, $\beta$ -galactosidase, β-glucosidase, β-glucuronidase, α-mannosidase and α-fucosidase activities; hydrolysis of aesculin, agar, casein, chitin, gelatin, starch and urea;
acid production from raffinose, D-ribose, D-xylose, sucrose, trehalose, D-cellobiose, N-acetylglucosamine, glycerol, inositol, D-sorbitol, sodium acetate, trisodium citrate, and D-mannitol; and production of H<sub>2</sub>S and indole.

The strains KMM 296 and *C. amphilecti* NRIC 815^T^ shared many common phenotypic features, such as the respiratory type of metabolism, motility by means of 1-2 polar and/or 2-3 lateral flagella, the ability to grow at 4-42°C, the presence of catalase, alkaline phosphatase, esterase (C4), esterase lipase (C8), leucine arylamidase, acid phosphatase, and α-glucosidase activities and assimilation of sucrose, maltose, sodium malonate, glycogen, D-mannitol, D-glucose, 3-hydroxybutyric acid, and L-proline (Table 2). They could not synthesize arginine dihydrolase, lipase (C14), cystine arylamidase, α-chymotrypsin, N-acetyl-β-glucosaminidase, β-glucosidase, α-galactosidase, β-glucuronidase, α-mannosidase, and α-fucosidase; hydrolyse agar, chitin, aesculin, gelatin, starch, urea, and Tween 80; produce acid from D-mannose, melibiose, raffinose, L-rhamnose, D-ribose, N-acetylglucosamine, inositol, D-sorbitol, glycerol, and D-mannitol; reduce nitrate to nitrite and assimilate L-arabinose, D-mannose, N-acetylglucosamine, adipate, phenylacetate, itaconic acid, sodium acetate, propionic acid, trisodium citrate, and 4-hydroxybenzoic acid. However, the strain KMM 296 can be distinguished from its closest phylogenetic relative by the several phenotypic traits, including the presence of cytochrome oxidase and the ability to assimilate capric and valeric acids, the inability to produce acid from a set of carbohydrates and to assimilate D-glucose, D-mannitol, maltose, D-gluconate, L-malate, L-rhamnose, N-acetylglucosamine, D-ribose, inositol, suberic acid, lactic acid, L-alanine, potassium 5-ketogluconate, 3-hydroxybenzoic acid, L-serine, salicin, melibiose, L-fucose, L-arabinose, L-histidine, and potassium 2-ketogluconate (Table 2). The above findings can extend the phenotypic characteristics those were reported for the species *C. amphilecti* by Romanenko et al. [3] after justification of the placement of strain KMM 296 in this species.

### The whole-cell fatty acid, polar lipid and respiratory quinone composition in the strains KMM 296 and *Cobetia amphilecti* NRIC 815^T^

The predominant fatty acids (>5% of the total fatty acids) of the strain KMM 296 were C_18:1_ ω7*c*, C_12:0_ 3-OH, C_18:1_ ω7*c*, C_12:0_ and C_17:0_ cyclo (Table 3).

**Table 3.** Fatty acid profiles (%) of the strains KMM 296 (1) and *Cobetia amphilecti* NRIC 815^T^ (2)

| Fatty acid* | 1 | 2 |
| --- | --- | --- |
| C<sub>10:0</sub> | 3.2 | 4.3 |
| C<sub>12:0</sub> | 11.0 | 8.7 |
| C<sub>16:0</sub> | 21.6 | 21.9 |
| C<sub>17:0</sub> cyclo | 5.2 | 9.4 |
| C<sub>16:1</sub> ω7c | 38.2 | 32.5 |
| C<sub>18:1</sub> ω7c | 14.9 | 12.7 |
| C<sub>19:1</sub> ω6c | 1.6 | tr |
| C<sub>12:0</sub> 3-OH | 15.8 | 20.3 |

\*Data are from the present study. Values are percentages of the total fatty acids. tr, Trace amount ( $\leq 1\%$ ).

The polar lipid profile of the strain studied was characterized by the presence of phosphatidylethanolamine, phosphatidylglycerol, phosphatidic acid, an unidentified aminolipid, an unidentified phospholipid and unidentified lipids, and it was found to be similar to that of *C. amphilecti* NRIC 815^T^ (Figure 1). The major respiratory quinone was Q-8, which is common among members of the class *Gammaproteobacteria*.

**Figure 1.**
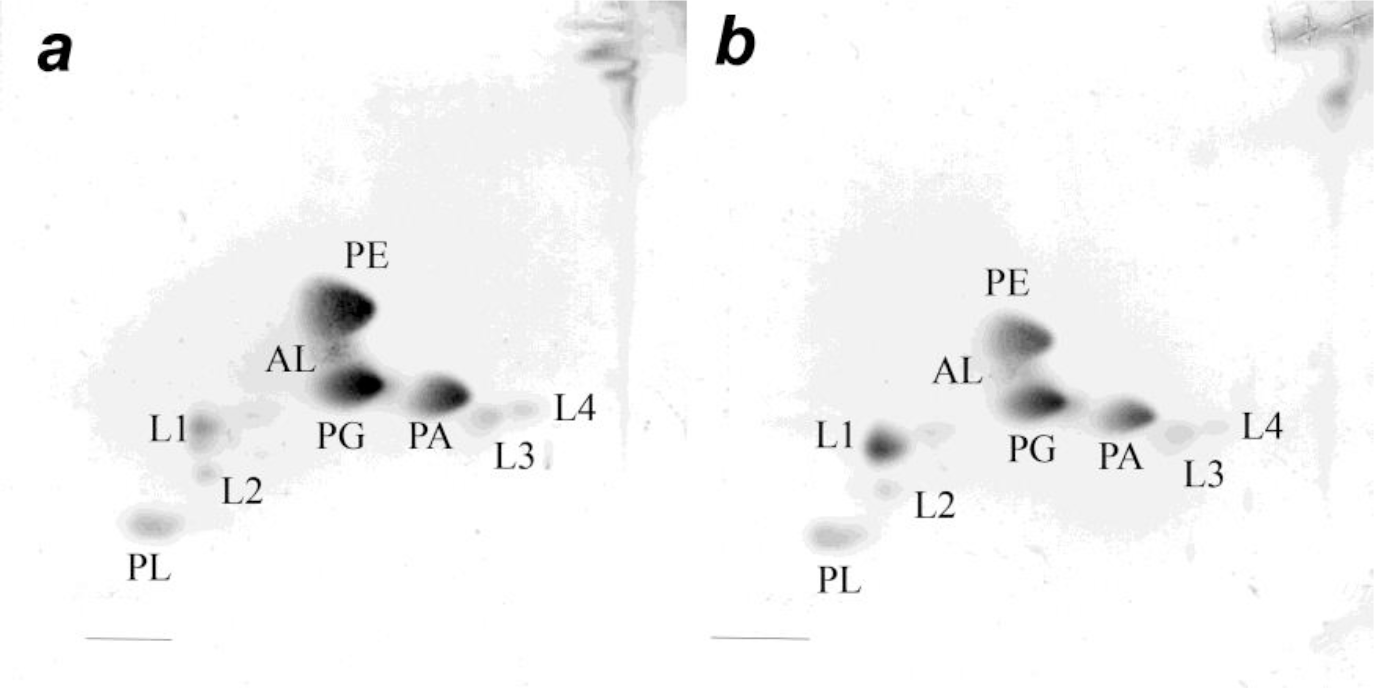
Two-dimensional thin-layer chromatogram of polar lipids extracted from strains KMM 296 (a) and *C. amphilecti* KMM 638^T^ (b). PE, phosphatidylethanolamine; PG, phosphatidylglycerol, PC, phosphatidylcholine, PA phosphatidic acid, AL, unidentified amino lipid; L, unidentified lipid, PL, unknown phospholipid

### Molecular phylogenetic analysis (16S rRNA) of the strains KMM 296, *Cobetia marina* LMG 2217^T^ and *Cobetia amphilecti* NRIC 815^T^

Phylogenetic analysis based on the 16S rRNA gene sequences revealed that the strain KMM 296 demonstrated only 99.5% sequence similarity to *C. marina* LMG 2217^T^ (=JCM 21022^T^). Whereas, it was found to be identical to the type strain of the other validly published species of the genus *Cobetia, C. amphilecti*, with 100% sequence similarity [3]. This suggests that the mussel isolate KMM 296 can be placed in this species instead of *C. marina*, as it was proposed previously [5]. In addition, the comparative genome analysis and phylogenomic analysis of the family *Halomonadaceae*, implemented by Tang et al. [29] indicated the significant differences between *C. marina* JCM 21022^T^ and KMM 296 (formerly named *C. marina* KMM 296) resulted from the sequence insertions or deletions and chromosomal recombination.

### Mol% G + C comparison between the strains KMM 296 and *Cobetia amphilecti* NRIC 815^T^

The genomic DNA G+C content of the strain KMM 296 was 62.5 mol% as determined by the genome sequencing data [9]. This value was slightly low than that obtained by the thermal denaturation method (62.7 mol%; [5]). The DNA–DNA relatedness between the mussel isolate and the strain *Cobetia amphilecti* NRIC 815^T^, which was determined by the experimental hybridization method, was 92%. This value was higher the 70 % threshold used for assigning bacterial strains to the same genomic species [30], and it strongly confirmed that the two strains belong to the single species *C. amphilecti*.

In addition, the closest evolutionary distances between the type strains of the species *C. amphilecti* and *C. litoralis*, on the one hand (99.93% 16S rRNA gene sequence similarity), and *C. marina* and *C. pacifica*, on the other hand (100% 16S rRNA gene sequence similarity), calculated by the EzBioCloud 16S RNA database tools and discussed earlier [31], assumed that the species *C. amphilecti* and *C. litoralis* belong to one species, and the species *C. marina* and *C. pacifica* could also be joined to a single species.

However, the comparison of the whole genome sequences of all *Cobetia* spp. strains, deposited currently in the NCBI GenBank database (Table 1), revealed that each genome contains up to seven 16S rRNA genes, with the different levels of the similarity (99,86-100%) within the strain, as well as between the species [31]. Therefore, the comprehensive whole genome-based analyses are required for the *Cobetia* species delineation.

### Whole genome-based phylogeny and analysis of *Cobetia* strains

In total, 36 *Cobetia* strains were chosen for phylogenetic and comparative analyses, eight of which have been sequenced by this study (the *Cobetia* spp. type strains: NRIC 0814^T^, NRIC 0815^T^; NRIC 0813^T^; the *Cobetia* spp. isolates 2AS1, 2AS, 1AS1, 29-18-1, 10Alg 146) and 28 were retrieved from the RefSeq database at NCBI. The genomic dataset included genomes of five type strains of five *Cobetia* species (as was stated previously [1-3], 9 strains of 4 species, and 22 *Cobetia* spp. strains. The overall features of the genomes are listed in Table 1. The genome size ranged from 3.1 to 4.6 Mbp with an average 4.1 Mbp, while the %GC content varies slightly and was 62−63%, except four strains with 57−57.5%. Apparently, such values might be due to the difference in the genome completeness levels. According to the NCBI Quality analysis (CheckM), the assemblies showed 81.26−99.65% completeness and 0.49−2.93% contamination (Table 1).

A core genome phylogeny was used to estimate the phylogenetic relationships of the *Cobetia* strains (Figure 2). According to the tree, the strains fall into three clades with subclades (bootstrap values = 100). The first one included three subclades and contained 19 strains, among which two type strains *C. litoralis* NRIC 0814^T^ and *C. amphilecti* NRIC 0815^T^ clustered at the same subclade. The second clade consisted of 13 strains, including the type strains *C. pacifica* NRIC 0813^T^ and *C. marina* JCM 21022^T^. The third one branched distantly and consisted of two *C. crustatorum* and two *Cobetia* spp. strains. According to the obtained topology of the phylogenomic tree, it is clear that five described species may actually represent three species (Figure 2).

**Figure 2.**
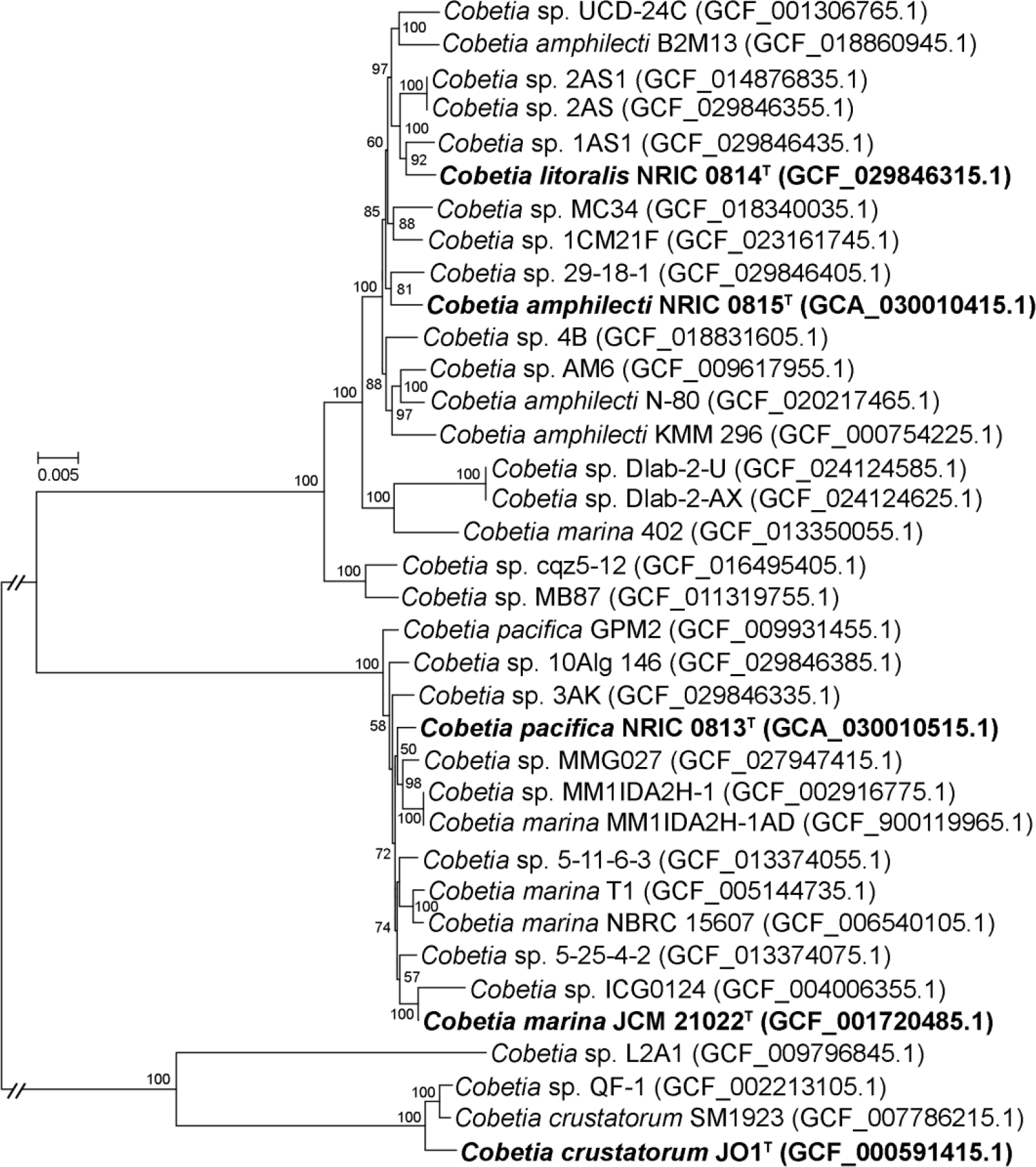
Maximum-likelihood phylogeny of the genus *Cobetia* based on concatenated 1432 single-copy core gene sequences and reconstructed by IQ-TREE with non-parametric bootstrapping using 100 replicates, including Bar, with 0.005 substitutions per amino acid position. The corresponding accession numbers for the genomes are given in parentheses. The type strains of *Cobetia* species are in bold.

The values of phylogenomic metrics ANIb, AAI, and dDDH are one more evidence to redefine the species assignments within the genus *Cobetia* (Supplementary Table S1). The obtained ANIb, AAI, and dDDH values among fourteen strains showing the same phylogenomic grouping, including the type strains *C. litoralis* NRIC 0814^T^ and *C. amphilecti* NRIC 0815^T^, were found to be 96.43−96.95%, 97.64−99.99% and 70.1−100%, respectively. The group of thirteen strains clustered together, including the type strains *C. pacifica* NRIC 0813^T^ and *C. marina* JCM 21022^T^, had showed the ranges 97.14−98.19%, 98.2−99.99%, and 80.4−100% of the ANIb, AAI, and dDDH values, respectively. These values between *C. crustatorum* JO1T, *C. crustatorum* SM1923, and *Cobetia* sp. QF-1 were 98.4−98.9%, 98.44−99.18%, and 89.5−91.7%, respectively. Considering the thresholds of 95-96% ANI, 95-96% AAI, and of 70% dDDH, defined for species demarcation, the type strains *C. litoralis* NRIC 0814^T^, *C. amphilecti* NRIC 0815^T^, *C. pacifica* NRIC 0813^T^, and *C. marina* JCM 21022^T^ do not belong to the corresponding originally assigned species [32]. The high values confirm phylogenetic grouping of those strains, which are likely represent two instead of four separate species. The phylogenomic metrics of the strains related to other clades were below the cutoff scores, implying that these might belong to several novel species of *Cobetia* (Table S1).

### Pangenome-based phylogeny and analysis of *Cobetia* strains

The pangenome of the genus *Cobetia* was performed to determine its genetic heterogeneity and phylogenetic relationships (Figure 3). The pangenome is presented by a set of the gene clusters (GCs) among which are the conserved core and accessory shell, and cloud genes. The core genes are found in ≥95% of the genomes, the shell genes are found more than 10% and less than 95% of the genomes, and the cloud genes presented in ≤10% of the genomes. Moreover, the single-copy genes (SCGs) as a part of the core found in all strains, while the unique genes (singletons) from the cloud are strain-specific. The pangenome of 36 strains of the genus *Cobetia* comprised a total of 6,648 gene clusters (Distance: Euclidean; Linkage: Ward) with 123,892 gene calls, that include 2,471 core gene clusters (93,289 genes in all 36 genomes), 1,469 gene clusters in the shell (26,722 genes), and 2,708 in the cloud (3,881 genes), including 1,902 gene clusters (1,920 genes) of singletons. It is interesting that 62 GCs were found exclusively to the strains grouped with the type strain *C. crustatorum* JO1^T^, while 20 GCs were found exclusively to the strains, clustered with the type strains *C. pacifica* NRIC 0813^T^ and *C. marina* JCM 21022^T^ (Figure 3). However, only one GC was found to be common for the 14 strains, grouped with *C. litoralis* NRIC 0814^T^ and *C. amphilecti* NRIC 0815^T^.

**Figure 3.**
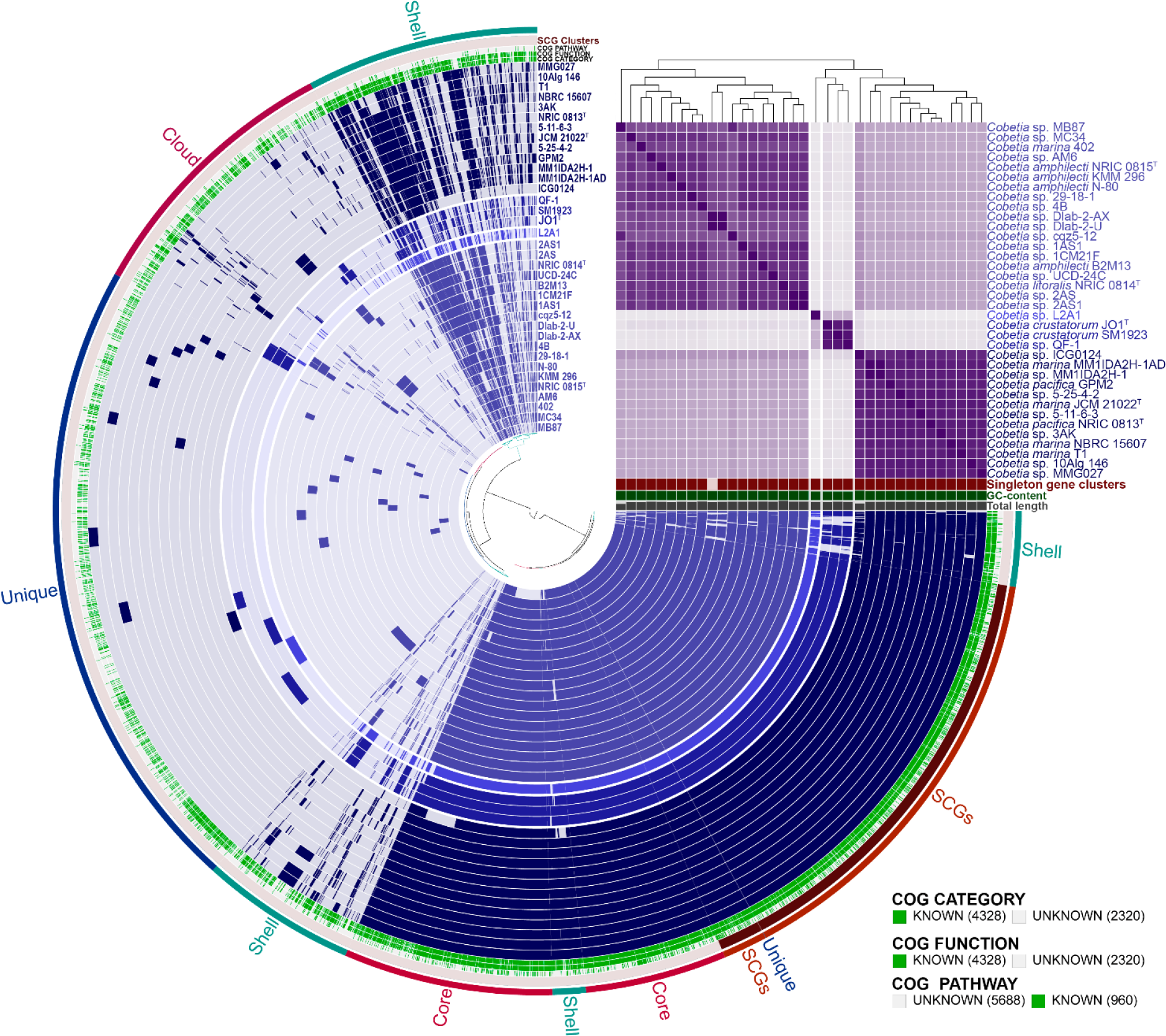
The pan-genome of 36 *Cobetia* spp. strains. Circle bars represent the presence/absence of 6,648 pan-genomic clusters in each genome. Gene clusters are organized as core, soft-core, shell, and cloud gene clusters using Euclidian distance and Ward ordination. The heatmap in the upper right corner displays pairwise values of average nucleotide identity (ANIm) in percentages

The core and unique gene clusters were further annotated into COG classes. The core genes were related to the following classes: translation and ribosomal biogenesis (10.48%), amino acid transport and metabolism (9.58%), cell envelope synthesis (7.35%), energy production and conversion (6.95%), transcription (6.65%), carbohydrate metabolism and transport (6.35%), lipid metabolism (6.06%), inorganic ion transport and metabolism (5.91%), coenzyme metabolism (5.46%), post translational modifications (5.21%), replication and repair (4.42%), signal transduction (3.82%). The classes for nucleotide metabolism and transport, secondary metabolite synthesis, defense mechanisms, cell cycle control, and intracellular trafficking and secretion were in a minority in the core (1.24–3.07%).

It is worth noting that each of *Cobetia* genome contains from one to 208 unique genes (Figure 4). The largest number of singletons was observed in the genomes of *Cobetia* sp. Dlab-2-U (208 genes), *C. crustatorum* SM1923 (164), *Cobetia* sp. L2A1 (144), and *Cobetia* sp. QF-1 (130). The genomes of *C. marina* MM1IDA2H-1AD, *Cobetia* sp. 2AS, *Cobetia* sp. 2AS1, and *Cobetia* sp. MM1IDA2H-1 account the smallest number of the unique genes 3, 3, 3, and 1, respectively. The remaining genomes harbors from 26 to 87 ones. According to the COG class annotation of these unique genes, the most abundant functional classes were cell wall/membrane/envelope biogenesis (14.12% of total unique gene clusters), general functional prediction only (10.63%), replication and repair (9.8%), defense mechanisms (6.98%), amino acid metabolism and transport (6.48%), and transcription (6.15%).

**Figure 4.**
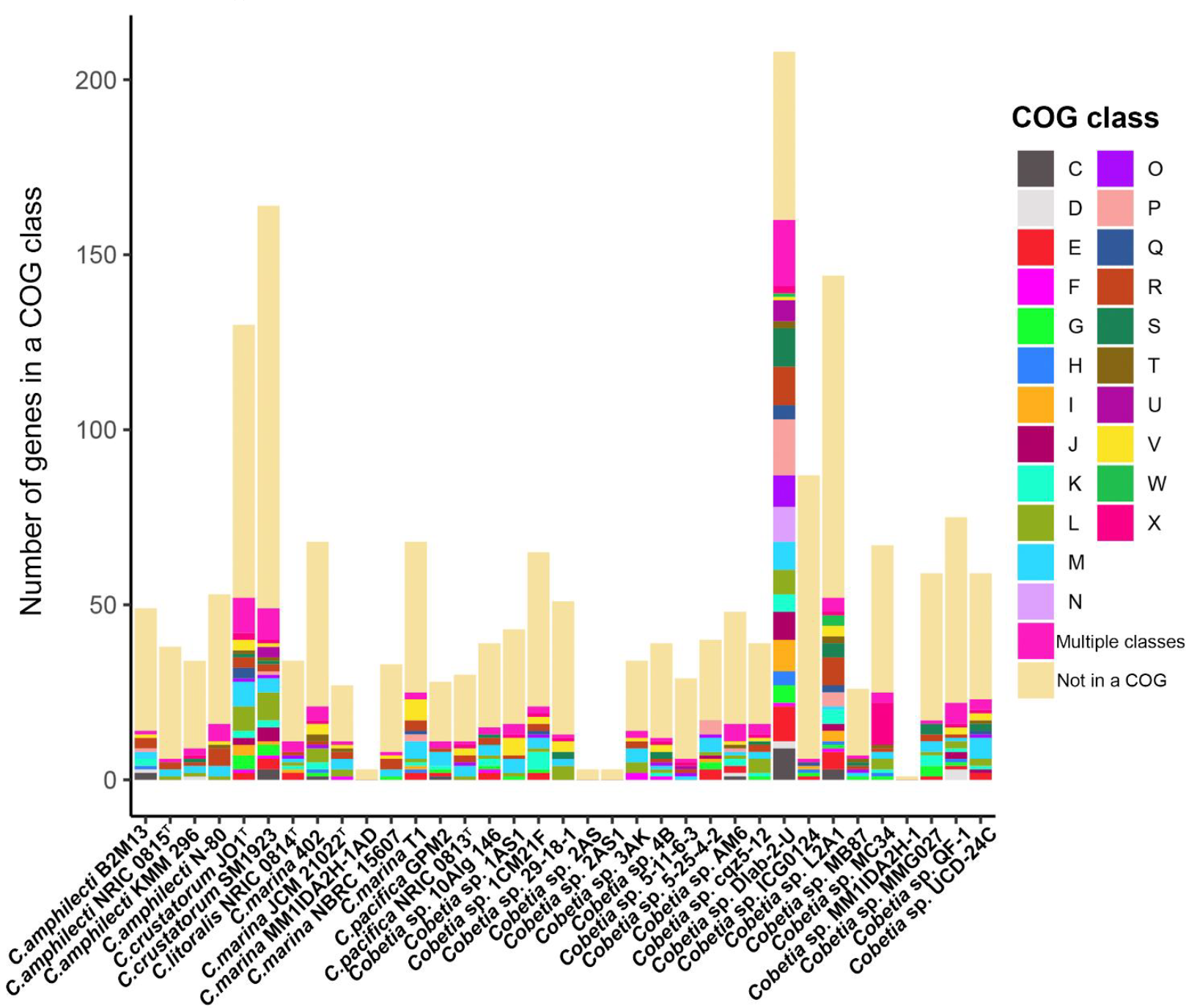
Number of unique genes assigned to a functional class (COG) among strains of *Cobetia* spp. Classes: C, energy production and conversion; D, cell cycle control and mitosis; E, amino acid metabolism and transport; F, nucleotide metabolism and transport; G, carbohydrate metabolism and transport; H, coenzyme metabolism, I, lipid metabolism; J, translation; K, transcription; L, replication and repair; M, cell wall/membrane/envelope biogenesis; N, cell motility; O, post-translational modification, protein turnover, chaperone functions; P, inorganic ion transport and metabolism; Q, secondary structure; R, general functional prediction only; S, function unknown; T, signal transduction; U, intracellular trafficking and secretion; V, defense mechanisms; W, extracellular structures; X, mobilome: prophages, transposons; Multiple classes, genes assigned to two or more COG categories; Not in a COG, COG not defined.

According to modeling of the pan- and core genome sizes upon the addition of new genomes into the dataset, the pangenome of *Cobetia* spp. is an open with a γ value of 0.43 (Figure 5 A). The best-fit regression curve for a pangenome is rising upwards, implying an expanding pangenome, while the core genome’s curve tends to reach a plateau. Moreover, the fitting of the curve to a power law showed that the number of the new gene cluster discovery with adding of new genome would add 52 and 29 genes to the pangenome as predicted for the 37^th^ and 100^th^ sequenced genomes, respectively (Figure 5 B).

**Figure 5.**
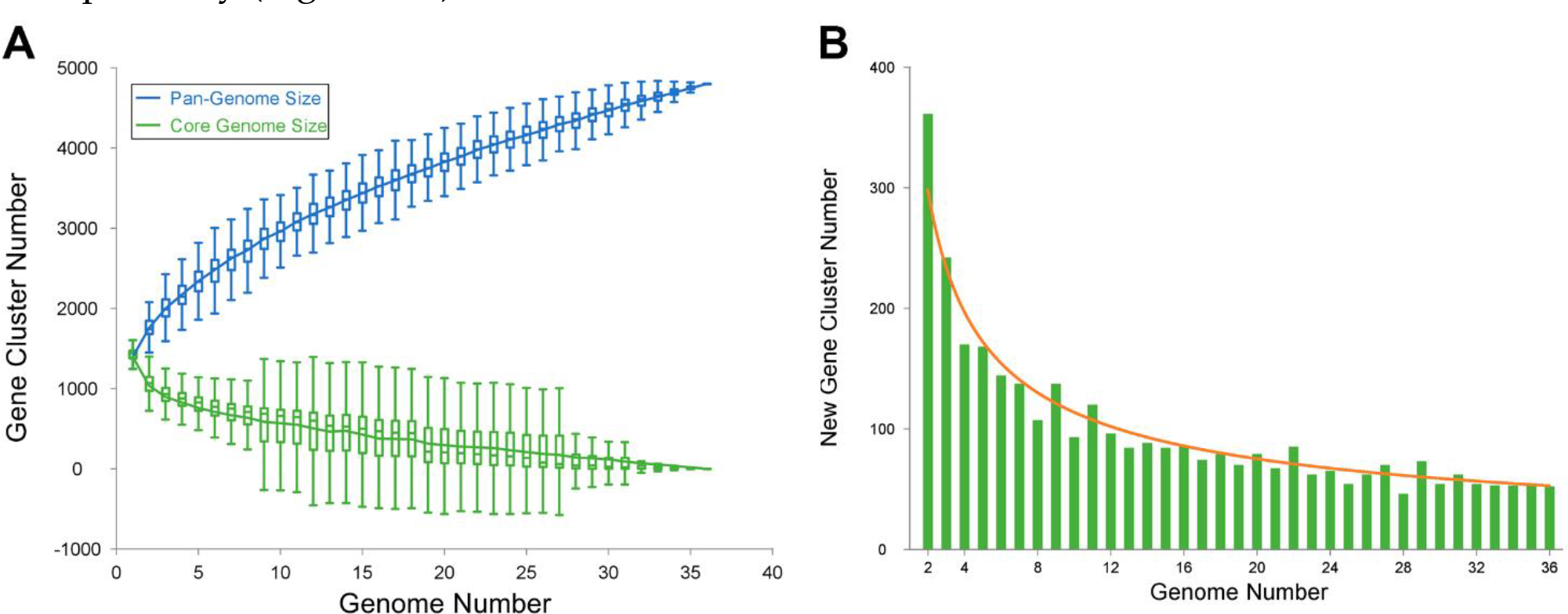
Pangenome modelling. (A) Gene accumulation curves of the pangenome and the core genome of 36 *Cobetia* genomes. Pangenome curve: *y*=889.06*x*^0.43^+535.51. Core genome curve: *y*=1250.45e^-0.07*x*^-29.77. (B) The new gene cluster number plot, curve: *y*=452.537*x*^-0.6^.

Therefore, based on the results of genomic, phylogenetic, phenotypic, and chemotaxonomic study, we suggested that the species *C. litoralis* should be placed in the species *C. amphilecti*, and the species *C. pacifica* should be included in the species *C. marina*. In accordance with the priority rules, *C. litoralis* should be reclassified as a later heterotypic synonym of *C. amphilecti*, and *C. pacifica* is a later heterotypic synonym of *C. marina*. The emended descriptions of the species *C. amphilecti* and *C. marina* are proposed.

### Emended description of the species *Cobetia amphilecti* Romanenko *et al*. 2013

The description of the species *Cobetia amphilecti* and *Cobetia litoralis* is as given by Romanenko *et al*. (2013), with the following amendments. Cells are oxidase-positive and motile by means of 1-2 polar and/or 2-5 lateral flagella. The predominant fatty acids (>5% of the total fatty acids) were C_16:1_ *ω*7*c*, C_12:0_ 3-OH, C_16:0,_ C_18:1_ *ω*7*c*, C_17:0_ cyclo and C_12:0_. The polar lipid profile is characterized by the presence of phosphatidylethanolamine, phosphatidylglycerol, phosphatidic acid, an unidentified aminolipid, the two unidentified phospholipid and the four unidentified lipids. The major respiratory quinone is Q-8. The genomic DNA G+C content is 62.5 mol%.

### Emended description of the species *Cobetia marina* (Cobet *et al*. 1970) Arahal *et al*. 2002 Romanenko *et al*. 2013

The description of the species *Cobetia marina* and *Cobetia pacifica* is as given by Arahal *et al*. (2002) and Romanenko *et al*. (2013), with the following amendments. Cells are oxidase-positive and motile by means of 1-2 polar and/or 2-5 lateral flagella. The predominant fatty acids (>5% of the total fatty acids) were C_16:1_ *ω*7*c*, C_12:0_ 3-OH, C_16:0,_ C_17:0_ cyclo, C_18:1_ *ω*7*c* and C_12:0_. The polar lipid profile is characterized by the presence of phosphatidylethanolamine, phosphatidylglycerol, phosphatidic acid, an unidentified aminolipid, the two unidentified phospholipid and the four unidentified lipids. The major respiratory quinone is Q-8. The genomic DNA G+C content is 62.2-62.4 mol%.

## Notes

### Competing Interest Statement

The authors have declared no competing interest.

